# Structural insights into the interaction of heme with protein tyrosine kinase JAK2

**DOI:** 10.1101/2020.08.13.246454

**Authors:** Benjamin Franz Schmalohr, Al-Hassan M. Mustafa, Oliver Krämer, Diana Imhof

## Abstract

Janus kinase 2 (JAK2) is the most important signal transducing tyrosine kinase in erythropoietic precursor cells. Its malfunction drives several myeloproliferative disorders. Heme is a small metal ion-carrying molecule, which is incorporated into hemoglobin in erythroid precursor cells to transport oxygen. In addition, heme is a signaling molecule and regulator of various biochemical processes. Here we show that heme exposure leads to hyperphosphorylation of JAK2 in a myeloid cancer cell line. Two peptides identified in JAK2 represent heme-regulatory motifs and show low micromolar affinities for heme. These peptides map to the kinase domain of JAK2, which is essential for downstream signaling. We suggest these motifs to be the interaction sites of heme with JAK2, which drive the heme-induced hyperphosphorylation. The results presented herein may facilitate the development of heme-related pharmacological tools to combat myeloproliferative disorders.

## Introduction

Janus kinase 2 (JAK2), a protein tyrosine kinase, is crucial for the transduction of cytokine and hormone signals in humans and other vertebrates (1). Defective JAK2 signaling is involved in myeloproliferative disorders such as polycythemia vera, leukemia, and lymphoma (2–5). Under physiological conditions, JAK2 activity is tightly controlled via autoinhibitory phosphorylation mediated by the Janus homology 2 (JH2) pseudokinase domain (6, 7). Its action is further regulated by cellular inhibitors, such as lymphocyte adapter protein (LNK), casitas B-lineage lymphoma protein (CBL), suppressor of cytokine signaling 3 (SOCS3), and protein tyrosine phosphatases (PTPs) (8–10). Pharmacological inhibitors against aberrant JAK2 activity have been developed recently (11).

In 2010, JAK2 was found to be regulated by heme (iron (II/III) protoporphyrin IX) (12). Heme is a known regulator of protein function and stability, controlling hemoglobin synthesis, its own biosynthesis, as well as inflammatory processes (12–17). Regulatory heme binding occurs primarily via heme-regulatory motifs (HRMs) (13) – short sequence stretches on the protein surface, which contain a central, iron-coordinating amino acid (14, 18). The most well-known HRM is the Cys-Pro dipeptide (CP) motif, which consists of cysteine and proline (19). However, we have recently identified potential new HRMs based on histidine and tyrosine, e.g. HXH, HXXXY, and HXXXH (20, 21). The concept of heme binding to JAK2 is conceivable since the cells in which it is expressed, such as erythroid precursor cells, exhibit high heme concentrations (22, 23). JAK2 activity is activated by erythropoietin, which drives hemoglobin synthesis (24). At the same time, heme is imported into erythroid precursors via feline leukemia virus subgroup C receptor-related protein 1 (FLVCR1) (25) and then increases globin production via inactivation of the transcription inhibitor Bach1 (BTB domain and CNC homolog 1) (26). The aforementioned JAK2-heme interaction was initially identified by a database search for CP-containing peptides (12), but no proof of heme binding to a CP motif in JAK2 exists as of yet. We used K562 cells to investigate the effect of heme on JAK2 and hypothesize that a histidine/tyrosine (H/Y)-based and a CP motif of the JH1 domain are involved in heme-induced JAK2 phosphorylation.

## Results

### JAK2 is hyperphosphorylated upon heme exposure

To investigate the effect of heme on JAK2 we utilized the erythroleukemia cell line K562. Under normal conditions, these cells have low intrinsic JAK2 signaling and therefore low amounts of Y^1007^/Y^1008^ phosphorylated JAK2. However, heme is ubiquitously present, so that cells and the cell culture additive fetal bovine serum (FBS) contain a pool of labile heme (27, 28). This naturally occurring heme can mask heme-induced effects and therefore needs to be reduced to minimal amounts (12). We blocked intrinsic heme synthesis with the inhibitor succinyl acetone (SA) and extrinsic heme by using heme-depleted FBS (28). Western blot analysis of cell lysates from K562 cells treated in this way showed minimal JAK2 phosphorylation (Fig. 1). Upon addition of 2 *μ*M and 4 *μ*M heme, respectively, JAK2 was strongly phosphorylated (Fig. 1). This effect did not increase further upon addition of 10 *μ*M heme (data not shown). NaOH (negative control), which was used to dissolve heme, did not show any effect. We thus demonstrate that heme has a marked activating effect on JAK2 in a human erythroleukemia cell line.

**Figure 1.**
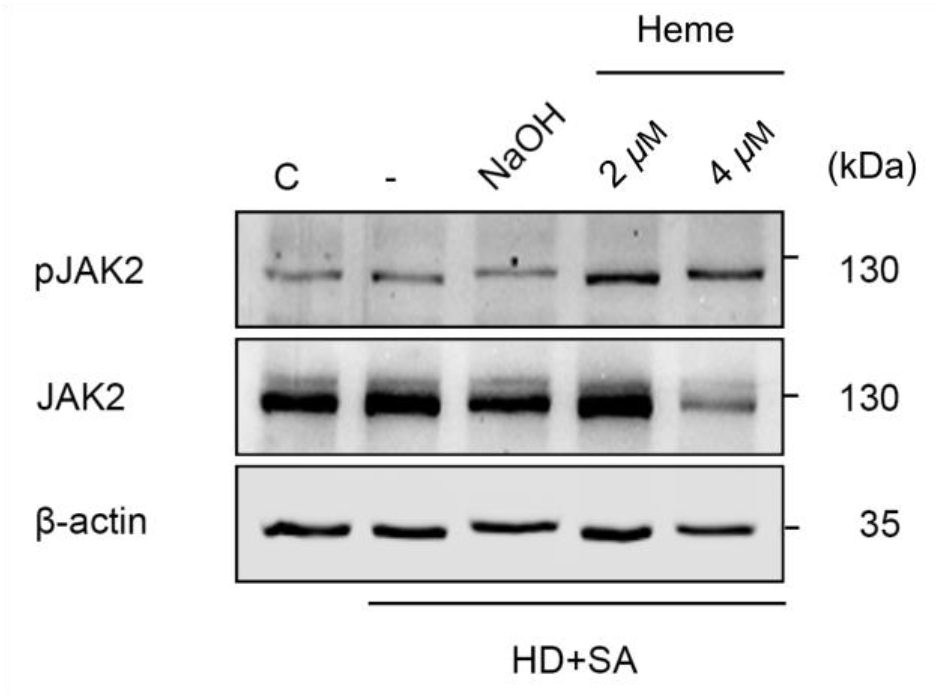
JAK2 is hyperphosphorylated upon heme exposure. K562 cells were depleted of heme by addition of succinyl acetone (SA) and heme-depleted medium (HD). Western blot analysis showed clear increase of JAK2 phosphorylation at Y^1007^/Y^1008^. NaOH did not increase JAK2 phosphorylation. The results are representative of three independent experiments.

### JAK2 possesses potential heme-regulatory motifs

We subsequently examined whether the heme effect was due to a direct heme-JAK2 interaction and if so, where the interaction site(s) could be located. Recent investigations revealed that His/Tyr-based motifs, such as (H/Y)X(H/Y), are interesting candidates for HRMs beside the CP motif (21). In a consensus sequence-based search (18), we identified the motif KR**Y**I**H**^974^RDLA (peptide 1) as potential HRM in the JH1 domain of JAK2 (21). This motif fulfills all criteria for regulatory heme binding (18, 29, 30). It has a positive net charge, stemming from basic amino acids, and contains advantageous hydrophobic amino acids. The central histidine, here H974, has been shown to effectively coordinate the heme iron ion, especially in the context of an H/Y motif such as YXH (20).

Previously, it was hypothesized that a CP motif might be responsible for heme binding to JAK2, but no proof of this has been brought forward yet (12). There is only a single CP motif in JAK2, RPDG**C**^1092^**P**DEI, which is also located within the JH1 domain (Fig. 2).

**Figure 2.**
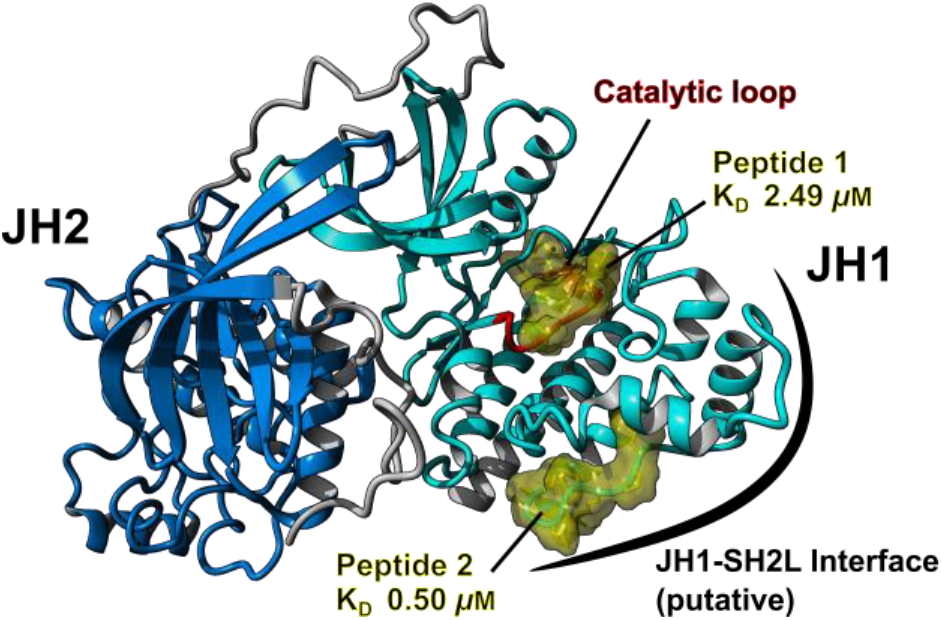
Visualization of the suggested motifs in the JH1 domain of JAK2. The motifs KR**Y**I**H**^974^RDLA (peptide 1) and RPDG**C**^1092^**P**DEI (peptide 2) are shown in yellow in the context of JH1 (light blue ribbons) and JH2 (dark blue ribbons). Peptide 1 is located in the catalytic loop (red ribbons), while peptide 2 is located in the putative JH1-SH2L interface (34). Visualization was performed using YASARA (39) and a model of JH1-JH2 (32).

Peptide 1, and the respective CP-containing peptide 2 were produced using solid-phase peptide synthesis and analyzed for heme binding using UV/Vis spectroscopy (21, 31). Both peptides were able to bind heme with good to high affinity, as apparent from the dissociation constants (KD) 2.49 ± 0.13 *μ*M (peptide 1) and 0.50 ± 0.23 *μ*M (peptide 2) (21, 31). The peptides form a 1:1 complex with heme as identified from a shift of the ν3-band to 1495 cm^−1^ (peptide 1) and 1491 cm^−1^ (peptide 2) in resonance Raman spectroscopy (21, 31). In consequence, both peptides represent suitable candidates for the heme interaction sites in JAK2.

### HRMs are localized in important regions of JAK2 JH1 domain

In order to visualize the heme-binding motifs, we used a JH1-JH2 domain model, which was generated from high-resolution crystal structures of JH1 and JH2 using extensive all-atom Molecular Dynamics (MD) simulations (32). The sequence corresponding to peptide 1 spans over the majority of the catalytic loop of the JH1 domain, which is located in the domains core (Fig. 2) (33). Parts of the motif are solvent-accessible, but heme binding might interfere with substrate binding at this site. Peptide 2, on the other hand, is located in a flexible loop on the surface of the JH1 domain, which has been hypothesized to be part of the JH1-Src-homology 2-like domain (SH2L) interface (34). Its surface accessibility would be beneficial for unhindered heme binding, as has been shown for other heme-regulated proteins (35).

It is also conceivable that further motifs are involved in heme binding to JAK2. The algorithm HeMoQuest (31) suggested 14 further motifs. However, the majority of these were predicted to possess a lower binding affinity compared to peptides 1 and 2 or are not favorably positioned on the surface of the protein.

## Discussion

Here we suggest regulation of JAK2 by heme in K562 cells, which is mediated by two potential heme-regulatory motifs. We confirmed these motifs on the peptide level and mapped their localization within the JAK2 structure to the catalytic loop and the putative JH1/SH2L domain interface. Our results are in good agreement with previous results by Zhang et al., which showed that JAK2 is heme-regulated in HeLa cells (12). In contrast to this report, however, we had a closer look at the possible heme interaction sites. Peptide 1 might not be favored for heme binding, since heme binding to the catalytic loop could interfere with phosphorylation, for example by blocking access to the catalytic center. In contrast, heme binding to peptide 2 might disrupt the JH1-SH2L interaction and thus release the JH1 domain into the putative elongated active state, which would explain the observed hyperphosphorylation (34).

Alternatively, one could speculate that the increased heme phosphorylation could stem from an indirect effect. Heme could either directly bind to proteins/peptides that are upstream of JAK2 (e.g. type I or type II receptors or their ligands) (36), or it could induce other pathways, which may eventually lead to JAK2 activation (e.g. the TLR4-NFκB axis via secretion of cytokines) (37). Future studies will shed light on the underlying mechanism, yet are dependent on the availability of the 120 kDa protein JAK2.

In conclusion, we confirm heme binding to JAK2 and pinpoint two possible binding sites as supported by peptides representing the respective motifs. This study provides deeper insight into the regulatory effect of heme and might aid in unraveling the role of heme in JAK2-related diseases. Especially in myeloid leukemia, the heme-degrading enzyme heme oxygenase 1 has recently been identified as druggable target (38). Consequently, localizing heme-binding sites may aid in the development of novel research tools and subsequently new targeted drugs.

## Experimental procedures

### Reagents

Endotoxin-free heme (Fe(III)protoporphyrin IX chloride) was purchased from Frontier Scientific, Logan, UT. Succinylacetone (4,6-Dioxoheptanoic acid) was purchased from Sigma Aldrich, St. Luis, MO. FBS was purchased from Thermo Fisher Scientific, Waltham, MA. Heme-depleted medium was prepared from heme-depleted FBS, as described earlier (28). K562 cells were cultured at 37 °C and a 5% CO2 humidified conditions in RPMI-1640 medium supplemented with 5%–10% heme-depleted FBS, and 1% penicillin/streptomycin (Sigma-Aldrich, Taufkirchen, Germany). For immunoblot analysis, anti-pERK1/2 (Tyr202/Tyr204) (#9101), anti-JAK2 (#3230), anti-rabbit IgG HRP-linked (#7074), anti-mouse IgG HRP-linked (#7076) were purchased from Cell Signaling Technology, Frankfurt, Germany; anti-pJAK2 (Tyr1007/1008) (sc-21870), and anti-β-actin (sc-47778) were purchased from Santa Cruz, Heidelberg, Germany.

### Heme addition to K562 cells and measurement of JAK2 phosphorylation

K562 cells were depleted of endogenous heme by incubation with HD medium and 0.5 mM SA for 24 hours prior to heme addition. Heme was dissolved in 30 mM NaOH to a concentration of 500 *μ*M for 30 min in the dark and sterilized using a 0.2 *μ*m filter. After filtration, the concentration was normalized using an extinction coefficient of 32.482 mM^−1^ cm^−1^ at 398 nm (21) and further diluted to in DPBS to 10x final assay concentration. Cells were seeded in serum-free medium with 0.5 mM SA in 6-well plates and the freshly prepared heme solutions were added. After incubation for 24 hours, the cells were harvested by centrifugation (300*g*/5 min). Cells were lysed in NET-N buffer (100 mM NaCl, 10 mM Tris-HCl pH 8, 1 mM EDTA, 10% glycerine, 0.5% NP-40; plus cOmplete protease inhibitor cocktail tablets (Roche) and phosphatase inhibitor cocktail 2 (Sigma)) for 30 min on ice, sonicated (10 s/20% amplitude) and centrifuged (18,800*g*/20 min/4 °C). Protein content of lysates was estimated via Bradford assay. Proteins were detected and analyzed by SDS-PAGE and western blotting using enhanced chemiluminescence on an iBright CL1000 imaging system (Invitrogen, Germany).

## Data availability

All data is contained within the manuscript.

## Acknowledgements

The authors would like to thank the group of Dr. David E. Shaw (D. E. Shaw Research, New York) for kindly providing the JAK2 JH1-JH2 model.

## Author contributions

All authors designed and planned the study. B.F.S. and A.M.M. designed the workflow, performed the experiments, and collected the data. All authors analyzed the data. The manuscript was written through the contribution of all authors.

## Funding and additional information

The authors are grateful for financial support by the German Academic Exchange Service (DAAD) to Al-Hassan M. Mustafa.

## Abbreviations

Bach1: BTB domain and CNC homolog 1
CBL: casitas B-lineage lymphoma protein
CP: cysteine-proline
FBS: fetal bovine serum
FERM: 4.1, ezrin, radixin, moesin
FLVCR1: feline leukemia virus subgroup C receptor-related protein 1
HD: heme-depleted
Heme: iron (II/III) protoporphyrin IX
HRM: heme-regulatory motif
H/Y: histidine and/or tyrosine
JH1/2: Janus homology 1/2
JAK2: Janus kinase 2
K_D_: dissociation constant
LNK: lymphocyte adapter protein
MD: Molecular Dynamics
PTP: protein tyrosine phosphatase
SA: succinylacetone (4,6-dioxoheptanoic acid)
SH2L: Src-homology 2-like
SOCS3: suppressor of cytokine signaling 3
UV/Vis: ultraviolet/visible.

